# Top-down design of protein nanomaterials with reinforcement learning

**DOI:** 10.1101/2022.09.25.509419

**Authors:** Isaac D. Lutz, Shunzhi Wang, Christoffer Norn, Andrew J. Borst, Yan Ting Zhao, Annie Dosey, Longxing Cao, Zhe Li, Minkyung Baek, Neil P. King, Hannele Ruohola-Baker, David Baker

## Abstract

The multisubunit protein assemblies that play critical roles in biology are the result of evolutionary selection for function of the entire assembly, and hence the subunits in structures such as icosahedral viral capsids often fit together with remarkable shape complementarity^1,2^. In contrast, the large multisubunit assemblies that have been created by *de novo* protein design, notably the icosahedral nanocages used in a new generation of potent vaccines^3–7^, have been built by first designing symmetric oligomers with cyclic symmetry and then assembling these into nanocages while keeping the internal structure fixed^8–14^, which results in more porous structures with less extensive shape matching between the components. Such hierarchical “bottom-up” design approaches have the advantage that one interface can be designed and validated in the context of the cyclic oligomer building block^15,16^, but the disadvantage that the structural and functional features of the assemblies are limited by the properties of the predesigned building blocks. To overcome this limitation, we set out to develop a “top-down” reinforcement learning based approach to protein nanomaterial design in which both the structures of the subunits and the interactions between them are built up coordinately in the context of the entire assembly. We developed a Monte Carlo tree search (MCTS) method^17,18^ which assembles protein monomer structures in the context of an overall architecture guided by a loss function which enables specification of any desired overall structural properties such as shape and porosity. We demonstrate the power of the approach by designing hyperstable icosahedral assemblies more compact than any previously observed protein icosahedral structure (designed or naturally occurring), that have very low porosity and are robust to fusion and display of proteins as complex as influenza hemagglutinin. CryoEM structures of two designs are very close to the computational design models. Our top-down reinforcement learning approach should enable the design of a wide variety of complex protein nanomaterials by direct optimization of overall system properties.

## Main Text

There has been considerable recent progress in protein nanomaterial design using a “bottom-up” hierarchical approach (Fig.1a, left) in which monomeric structures are first docked into symmetric oligomers^19–21^, and then these are assembled into closed assemblies with tetrahedral, octahedral or icosahedral symmetry^8,22,23,9–11,13,14^, or open assemblies such as 2D layers and 3D crystals^24–28^. An advantage of this hierarchical approach is that the multiple interfaces that stabilize the assembly can be validated independently (the first by characterization of the symmetric oligomer, the second, by characterization of the nanomaterial assembly from the preformed oligomer), considerably increasing the robustness of the overall design process. While such designed assemblies are already proving useful in biomedicine in immunobiology and other areas, as highlighted by the recent approval of a *de novo* designed COVID vaccine^5,29–31^, the bottom-up approach does have limitations: the properties of the assembly are limited to what can be generated from the available oligomeric building blocks, and more generally, at least one of the subunit-subunit interfaces must be strong enough to stabilize a cyclic oligomeric substructure in isolation. Because of these limitations, it has been challenging to directly optimize properties of the overall assembly; for example for cellular delivery applications non-porous assemblies analogous to icosahedral viral capsids could be very useful^32–35^, but the almost perfect fit between asymmetric building blocks in such structures is difficult to achieve using bottom-up approaches.

**Figure 1.**
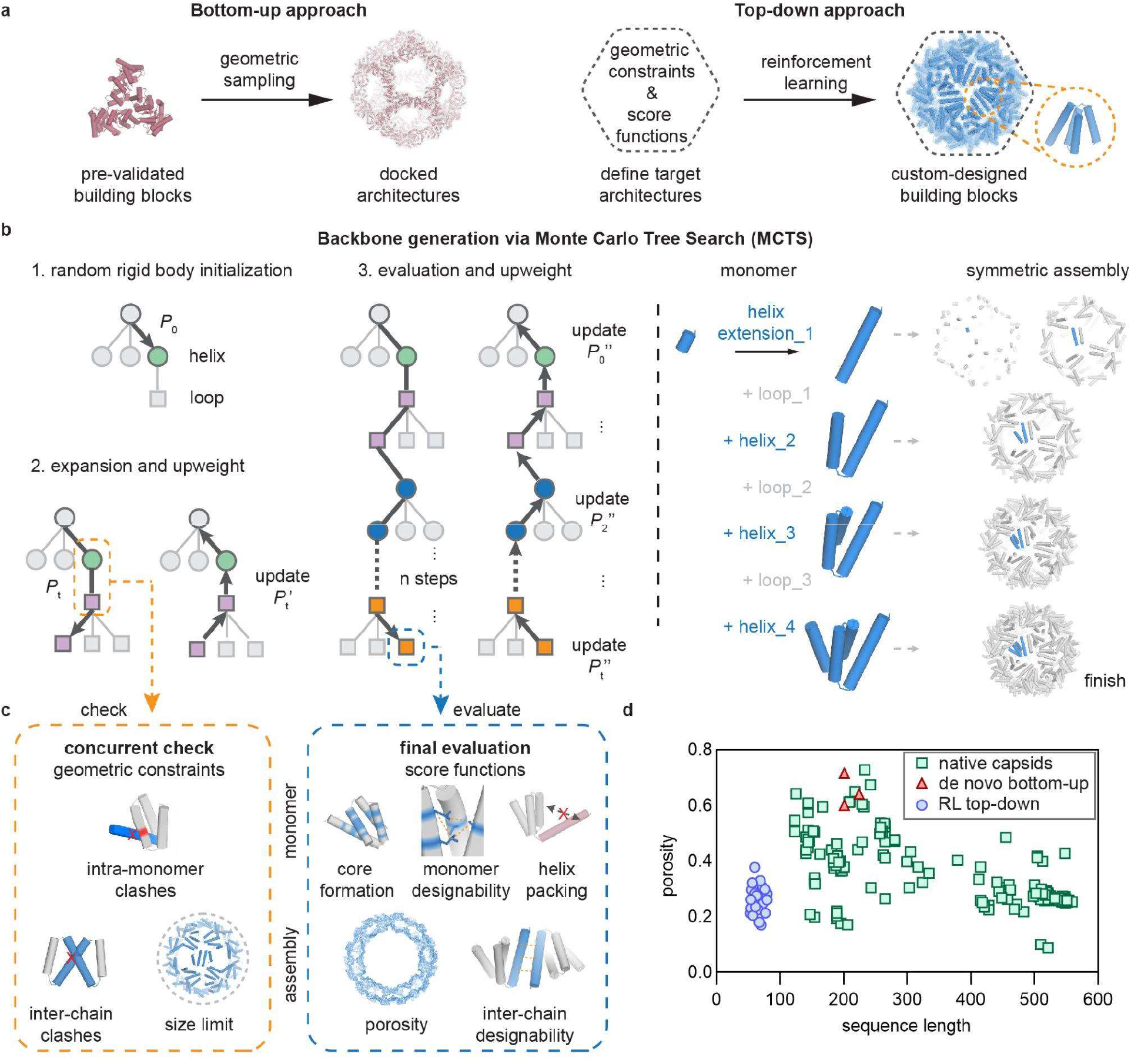
Top-down design strategy and computational pipeline.

**a**, Schematic illustrations of designing symmetric protein assemblies with bottom-up (left) and top-down strategies (right). **b**, Monte Carlo Tree Search (MCTS) architecture for monomer backbone generation. During each simulation, a helix stub is initialized at an random rigid body start position, different configurations of helices and loops were sampled and constructed sequentially with probability Pt stored in each edge to build the search tree. Each move was checked against a set of pre-defined geometric constraints during the expansion stage and update probabilities P_t_’ afterwards. Upon successful completion of a search tree, the monomer was scored by score functions and probabilities Pt” were back-propagated to update all the search tree edges. Symmetric transformations were applied to build an icosahedral capsid in parallel with monomers using the MCTS generative algorithm (right). **c**, Concurrent geometric check (left) is performed at every step of the expansion stage and violations would result in termination of the search tree; final evaluation (right) with a series of score functions is performed upon completion of a simulation for monomers and assemblies. **d**, RL-generated capsids occupy a distinct structural space as compared to de novo designed protein cages and natural capsids.

We sought to overcome the limitations of bottom-up protein nanomaterials design approaches by developing a top-down approach (Fig.1a, right) which starts from a specification of the desired properties (overall symmetry, porosity, etc) of the nanomaterial and systematically builds up subunits which pack together to optimize these properties. We reasoned that protein fragment assembly^36–40^, which can generate a very wide array of monomeric protein structures, could provide a suitable mechanism for generating diversity. Previous design approaches such as SEWING have built up proteins from fragments optimizing for monomer stability at each step^41^, but we aimed instead to optimize for overall system properties, which could involve trading off monomer stability for increased subunit-subunit interaction strength and other properties. To enable such end-state based optimization, we turned to reinforcement learning (RL), which has achieved considerable success recently in different fields of artificial intelligence, such as self-driving cars^42^ and the AlphaGo program that defeats top human players in the game of Go^43,44^. Monte Carlo tree search (MCTS)^17,18^ is an RL algorithm that finds optimal series of choices within a search tree. In MCTS, choices are selected randomly at each branch point to find a path down the tree, and after exploring a path, the state is evaluated, and probabilities at each branch point backpropagated up the tree are reweighted accordingly such that subsequent iterations are more likely to lead to optimal paths.

We sought to develop a MCTS algorithm for generating protein nanomaterials which builds up the monomeric subunits from protein fragments directly optimizing for prespecified global material properties. We set up the tree search such that at each step in the tree, a short protein fragment is appended at either the N-terminus or C-terminus of the growing chain. The number of fragments to consider at each step is a tradeoff between the rapidity of learning (with a smaller number, weights on each choice can be learned more quickly) and the total diversity of structures which can be generated (which increases with the number of choices at each step). We chose to balance these by using as building blocks parametrically generated straight helices, which are fully described by a single parameter (the length, which we allow to vary from 9 to 22 residues), followed by short loops clustered into 316 bins (derived from clustering loops in a large helical protein database, see Methods). The search begins with selection of one of the helix possibilities, and then alternates between addition of a loop or a helix choice at either terminus; once a loop bin is chosen, we select randomly from the closely related loop backbones within the cluster (Fig.1b, left). While this is a far narrower set of local structures than that observed in native protein structures, we found in preliminary explorations that a wide variety of compact protein shapes could be readily generated from such building blocks. Building up a 100 residue protein backbone with this approach requires on the order of 5 helix and 4 loop additions, yielding a total number of possibilities on the order of 1 × 10^17^ with additional structural diversity from the variation in loop backbones within a bin.

The search tree is modulated based on the specific problem specification at two levels: geometric constraints and score functions. First, potential moves consisting of helix or loop fragments are selected at each level of the search tree only if they pass geometric constraints that can be evaluated prior to assembly of the entire structure; these include internal clashes and overall shape constraints (see supplementary methods for full list of geometric constraints). Upon selection of a move passing the geometric constraints, its probability is upweighted, as are the probabilities of all prior moves leading to this point in the search tree. Second, completed backbones are evaluated using score functions that assess how well the overall generated structure satisfies the user specification of the problem to be solved (Fig.1c, supplementary methods), and the probabilities of selection of each possible move at each step along the search tree are reweighted accordingly. As individual move weights become increasingly biased after many traversals through the search tree, the generated complete backbones have higher and higher scores (Supplementary Fig.2). As each iteration takes on average only tens of milliseconds, high scoring backbones can be sampled at scale by searching over tens of thousands of iterations. To address the classical RL problem of balancing exploration with exploitation^42–44^, the search is initialized from many independent trees and the maximum probability of any one move is capped (supplementary methods).

We first tested the MCTS approach at the protein monomer level, choosing as a test problem the generation of protein backbones with arbitrarily prespecified overall shapes; to our knowledge, there are no current approaches for addressing this problem. A specified build volume is represented on a grid, and the MCTS is initialized randomly within the volume; at each move only additions that stay within the specified volume are accepted. For a range of prescribed shapes including regular polyhedra and letters from the alphabet, the ensembles of generated structures closely fill the specified volumes, and individual backbones have the prespecified shapes (Supplementary Fig.1). The average sequence length of the solutions increases through the optimization, as the process learns the choices of moves and combinations of moves that lead to satisfaction of the input constraints (Supplementary Fig.2a).

We next sought to generalize the MCTS to the design of symmetric nanomaterials by applying symmetry operators to generate assemblies with the desired symmetry at each step in the search tree. Each move (helix or loop addition) is assessed considering not only the growing monomer but also its interactions with all nearby symmetry mates; moves that introduce steric clashes are discarded (Fig.1b,c). We used this approach to build up icosahedral assemblies by using 59 transformation matrices to compute symmetry mates for a growing monomer. End-state based assembly score functions guide move probability reweighting at each step in the search tree; these include measures of cage porosity, interface designability, and to enable fusion constructs, external placement of at least one terminus (Fig.1c). We initialized millions of MCTS trajectories starting from a short helical fragment randomly placed within a specified upper distance bound of the origin in a random orientation, and for each carried out ten thousand iterations to generate a large set of diverse structures. The MCTS generated closely packed icosahedral assemblies which span a structural space distinct from that of native and previous *de novo* icosahedra, with shorter sequence lengths and lower porosities than any previously described protein icosahedra and comparable with the densely packed capsids generated by evolution (Fig. 1d).

The MCTS method rapidly generates tens of thousands of candidate icosahedral assemblies, and we experimented with approaches for rapidly designing sequences which stabilize these assemblies compatible with our overall top-down approach. In previous bottom-up nanocage design studies, the sequences and backbones of the oligomeric building blocks are pre-optimized, so only the new interface formed between the building blocks in the cage is designed, and the overall backbone is kept largely fixed^10^. In contrast, with the top-down MCTS approach, the entire sequence must be designed, with backbone relaxation to optimize sequence-structure compatibility both within and between the monomers and increase interface shape complementarity. A deep neural network trained to learn the sequence and structure relationships of native proteins was used to generate amino sequence profiles for each position in the newly generated backbones, which were used in turn to bias amino acid selection in the sequence design stage using Rosetta Design (see Methods). The resulting designs were filtered based on interface contact molecular surface area^45^, shape complementarity, predicted binding energy, exposed surface hydrophobicity, and AlphaFold (AF)^46^ prediction similarity to the design model (Methods). The rigid body and internal degrees of freedom of the selected icosahedral assemblies were then optimized by Rosetta symmetric relaxation^47,48^, starting from both the Rosetta design model of the assembly and the AF predicted structure of the monomer mapped back on to the assembly. To further increase sequence-structure compatibility, this design-predict-relax cycle was repeated three times, at each iteration carrying out sequence design on the full assemblies generated in the previous iteration, mapping back the predicted monomer structures into the assemblies, and relaxing the full structure in Rosetta. We applied this sequence design and backbone refinement procedure to 220,000 of the MCTS-generated backbones, and selected 350 designs for experimental characterization.

Linear gene fragments encoding each design with hexahistidine purification tags were cloned into an E. coli expression vector, and the proteins produced in E. coli in a 96-well format were purified by immobilized metal affinity chromatography (IMAC) pull-down. About 40% of the 350 designs were expressed and soluble as assessed by SDS-PAGE. To evaluate particle formation, we performed negative stain electron microscopy (nsEM) on the IMAC elution fraction for each soluble sample. Two designs (RC_I_1 and RC_I_2, RL capsid with I symmetry, design#1 and 2) formed uniform particles at the expected size and shape (Fig.2a,b). We scaled up expression and purification of these two designs, with size exclusion chromatography (SEC) of both samples yielding single peaks with an apparent molecular weight in the range expected for these icosahedral assemblies (Fig.2c,d). Small-angle x-ray scattering (SAXS) on the SEC-purified samples indicated that both designs form assemblies similar to the intended 3D configuration in solution (Fig.2e,g). The designed assemblies had the expected alpha helical circular dichroism (CD) spectra and apparent melting temperatures above 65 °C. Assembly morphologies were retained after 1 h treatment at 95 °C and with subsequent cooling to 25 °C, as confirmed by nsEM analysis (Fig.2f,h, and Supplementary Fig.11).

**Figure 2.**
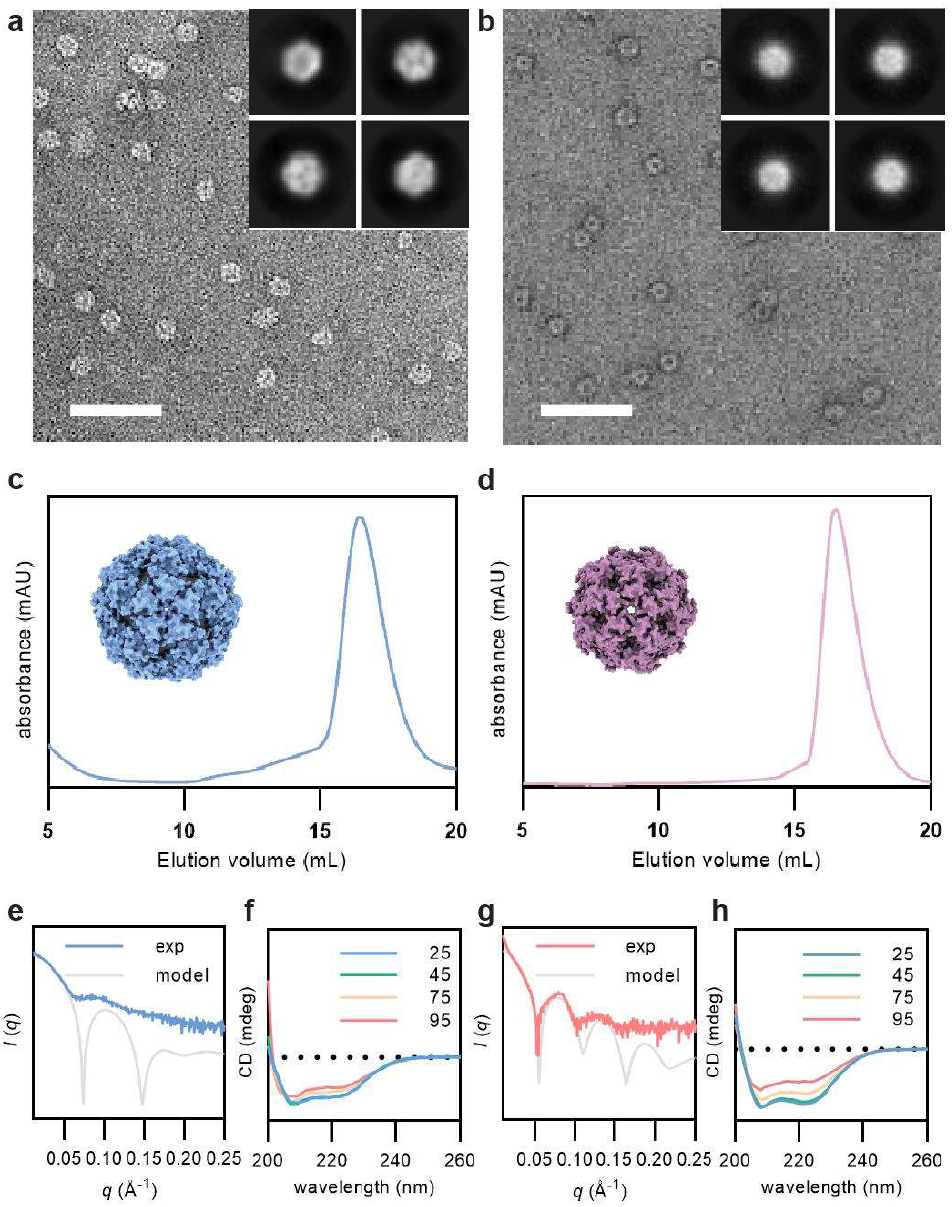
Experimental characterization of designed capsids RC_I_1 and RC_I_2.

**a,b**, Representative negative-stain EM micrographs and reference-free 2D class averages (inset) for RC_I_1 (left) and RC_I_2 (right), scale bar = 200 nm. **c,f**, A single peak was observed for each SEC elution profile near the expected elution volumes for the target complexes. Design models are shown as inset. **e,g**, 1D small-angle x-ray scattering patterns (SAXS, solid curves) of capsids in buffer solution match well with profiles calculated from the design models (gray dashed curves). **f,h**, Circular dichroism spectra measured at different temperatures showing designed capsids are highly thermally stable.

To evaluate the accuracy of our design strategy, we determined the structures of SEC purified RC_I_1 and RC_I_2 capsid particles using cryo-electron microscopy (cryo-EM, Fig.3). For RC_I_1, 3D reconstruction yielded a 2.5 Å resolution cryoEM atomic model that closely matched the computational design (Fig.3a-b). The N terminal helices of two monomers pack in an antiparallel fashion to form the C2-interface consisting of primarily nonpolar interacting side chains, while the 2-helices near C-terminus form the C5 interface with their neighbors (Fig.3b-c). Small apertures (diameter ~ 13 Å) present at the C3 axes of the capsid make the N-termini available for genetic fusion (Fig.3c). Over the designed monomer, the RMSD between cryoEM structure and design model is 0.76 Å (Fig.3d); a single rotamer flip (Phe63) and tilting of the C-terminal helix results in a slight expansion of the overall cage diameter resulting in an RMSD over all 60 subunits of 3.72 Å (Fig.3e). For RC_I_2, the 2.9 Å cryoEM structure of design RC_I_2 was even closer to the design model (Fig.3f-g), with RMSDs at the C2 and C5 interfaces of 0.66 and 0.27 Å, respectively (Fig.3h). The RC_I_2 monomer adopts the designed 3 helical bundle fold with a 0.59 Å RMSD to the design model (Fig. 3i), and the overall assembly is almost identical to the design model with a 1.39 Å RMSD over all 60 subunits (Fig.3j). The C2 interface is situated near the extended C-terminus of the monomer, allowing for potential monomeric or dimeric genetic fusions. The C5 pentameric interface is mediated by interactions between the N terminal helices, which point inward and enable functionalization of the interior of the capsid. With diameters of 13 and 10 nm for RC_I_1 and RC_I_2, respectively, and associated monomer lengths of 67 and 54 residues, the designed mini capsids are smaller than viral capsids, and to our knowledge, represent the smallest designed icosahedral biomolecular nanostructures to date.

**Figure 3.**
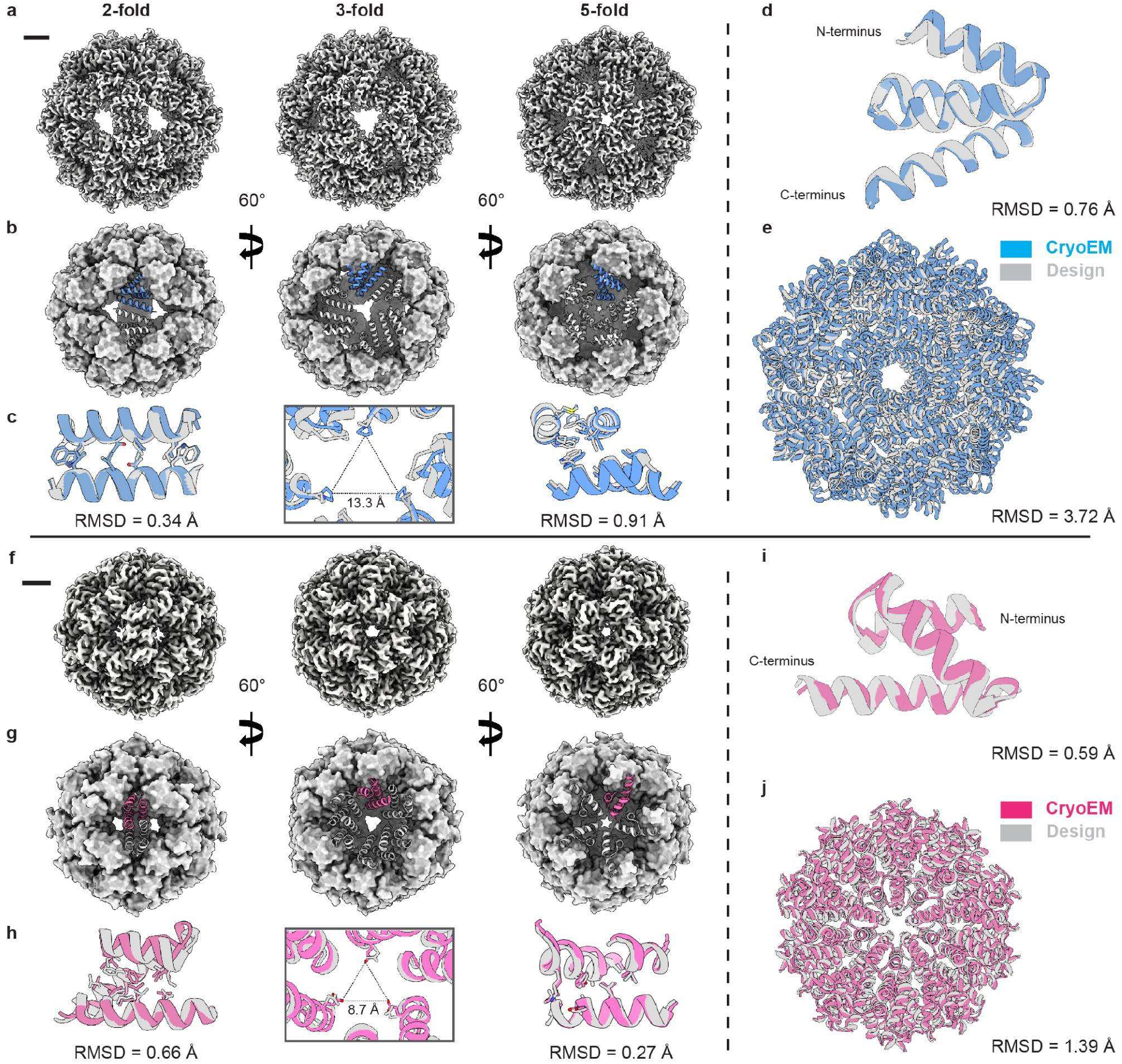
Near-atomic resolution cryoEM structures of designed capsids match design models. **a**, 2.5 Å cryoEM reconstruction of RC_I_1 viewed along the three symmetry axes (scale bar = 20 Å). **b**, CryoEM structure of RC_I_1 highlighting monomer packing and interfaces along each symmetry axis. **c**, Overlay and RMSD calculations for RC_I_1 as compared to the design model for each symmetry interface (cryoEM = blue; design = grey). **d**, Overlay and rmsd calculation for a single monomer of RC_I_1. **e**, Overlay and rmsd calculation for the entire 60-mer RC_I_1 capsid. **f**, 2.9 Å cryoEM reconstruction of RC_I_2 viewed along the three symmetry axes (scale bar = 20 Å). **g**, CryoEM structure of RC_I_2 highlighting monomer packing and interfaces along each symmetry axis. **h**, Overlay and rmsd calculations for RC_I_2 as compared to the design model for each symmetry interface (cryoEM = blue; design = grey). **i**, Overlay and rmsd calculation for a single monomer of RC_I_2. **j**, Overlay and rmsd calculation for the entire RC_I_2 capsid.

### Applications of designed capsids

The compact size and corresponding small exterior surface area of the designed particles enables the display of 60 or 120 copies of N and/or C terminal fused proteins with exceptionally high density, six or more times higher than previously designed icosahedral cages. As a first step to exploring applications of these structures, we used a recently developed sequence design method, ProteinMPNN^49^, which has been found to increase the robustness of protein designs, to generate 12 sequences for the RC_I_1 capsid backbone and filtered using the AF and Rosetta metrics described above. Two designs, RC_I_1-H9 and RC_I_1-H11 (the former designed by ProteinMPNN using the working capsid backbone Cα coordinates as input, the latter, the idealized polyA backbone without any backbone optimization and relaxation), assembled into the designed I1 symmetric capsid as evidenced by IMAC, SEC, and nsEM (Supplementary Fig.12-13). A 3 Å cryoEM structure of RC_I_1-H11 was almost identical to the design model, with a monomeric RMSD of 0.60 Å (Fig.4a), and a remarkably low full-cage RMSD over all 60 subunits of only 0.96 Å. RC_I_1-H9 and RC_I_1-H11 have on average 46 % sequence difference as compared to the parent capsid, and 30 % sequence difference with each other, including highly diverse interface residue selections (Fig.4a; for example the errant Phe63 of the parent capsid was redesigned to Glu63 in RC_I_1-H11, likely accounting at least in part for the closer agreement of RC_I_1-11 with the design model). These results also demonstrate that the RL approach can generate directly designable protein backbone geometries with a high degree of accuracy.

**Figure 4.**
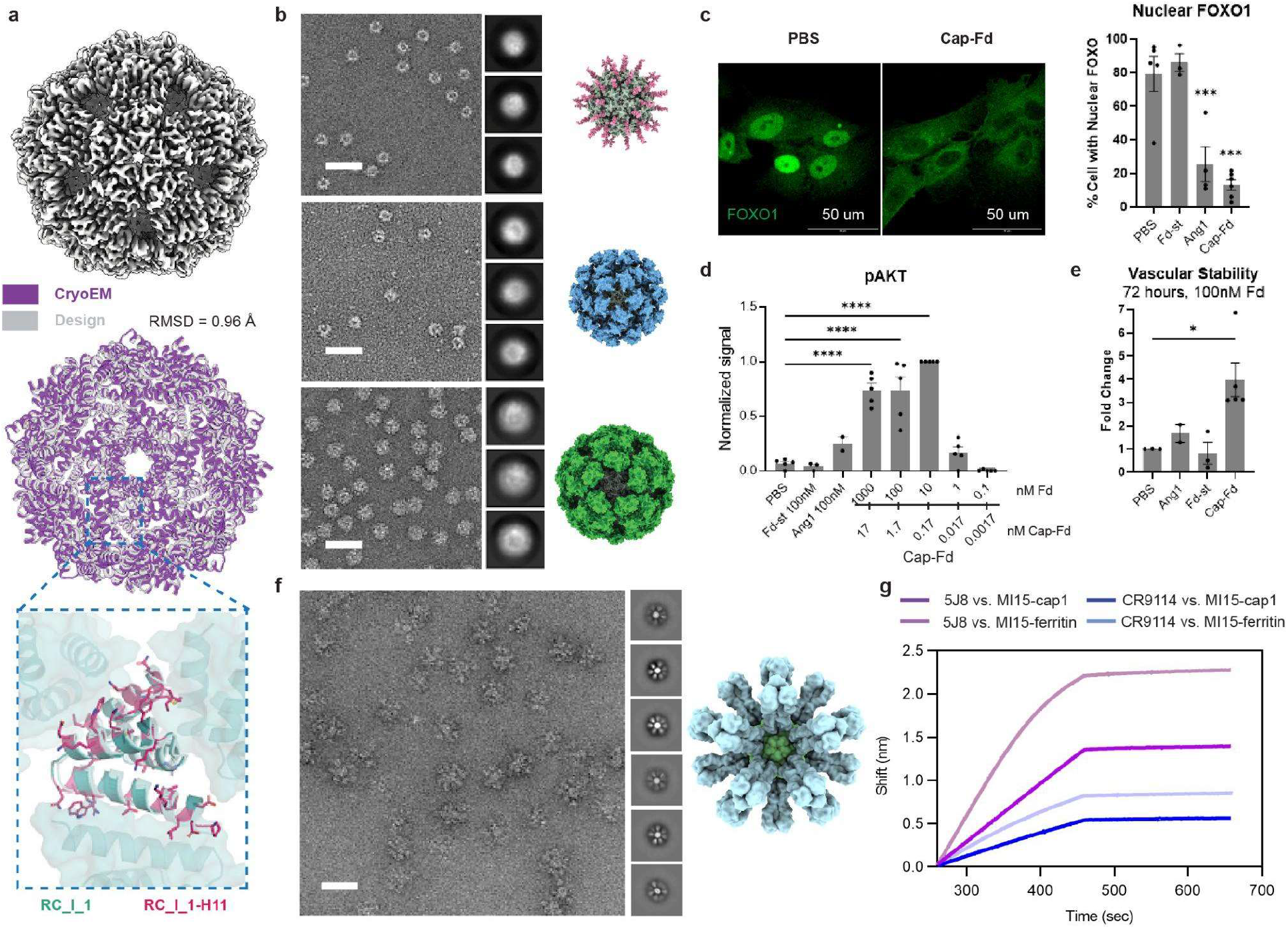
Applications of designed capsids.

**a**, Robustness of RC_1_1 capsid to sequence redesign using ProteinMPNN. The 3D cryoEM map at 3 Å resolution (top) and superposition of the structure on the design model (rmsd = 0.96 Å) at 3 Å resolution show that the RC_I_1-H11 structure is nearly identical to RC_I_1 despite considerable sequence differences (bottom; residue differences are highlighted in red (RC_I_1-H11) and teal (RC_I_1). **b**, From top to bottom, representative nsEM images and models of spyTag-, spyCatcher-, and GFP-fused (to N-terminus) RC_I_1-H11 with 2D class averages, scale bar = 50 nm. **c**, Capsid-Fd activates Tie2 downstream Akt phosphorylation and FOXO1 translocation. Serum starved HUVECs were treated with serially diluted Cap-Fd (1000-0.1 nM), Fd-st (100 nM), Ang1 (100 nM), or PBS control for 15 minutes before protein lysate collection for western blot analysis or PFA fixing for FOXO1 antibody stain. Left, Representative confocal images of HUVECs immunofluorescent stained with FOXO1 antibody after Cap-Fd (100nM) administration or PBS. Levels of significance were compared to PBS control. Right, Quantification showing percent of cells with nuclear FOXO1, 100 cells per counted in each biological replicate. **d**, Quantification of western blot showing pAKT signal normalized to Cap-Fd at 10 nM. P-values were calculated using One-way ANOVA with Bonferroni’s multiple comparisons test in Prism, Graph Pad. **e**, Quantification showing the vascular stability by averaging the number of nodes, meshes, and tubes calculated at 72 h timepoint using Angiogenesis Analyzer plugin in ImageJ. P-values were calculated using One-way ANOVA in Prism, Graph Pad. **f**, (right) Design model of 20 trimeric hemagglutinin proteins (HA) fused to a single RC_I_1 particle. (left) Representative nsEM micrograph and 2D class averages (inset) of mammalian cell secreted HA-capsids, scale bar = 50 nm. **g**, Bio-Layer Interferometry (BLI) data shows neutralizing antibody binding to HA-capsids comparable to canonical HA-ferritin constructs.

We next evaluated the robustness of the designs to genetic fusion by fusing spyTag, spycatcher, and GFP proteins to the RC_I_1-H11 capsid with a N-terminal (GGS)n linker (Fig.4b). In all cases, SEC elution profiles and nsEM micrographs showed monodisperse particles of the expected size and shape (Supplementary Information). 2D class averages (inset) revealed spherical structures similar to that of the original icosahedral capsid, with additional density at the periphery of particles, consistent with fused proteins connected to scaffolds via a flexible linker.

To assess the efficacy of the designed capsids in activating cellular signaling pathways by clustering cell surface receptors, we fused 60 copies of the angiopoietin 1 F domain, which binds the Tie2 receptor, to RC_1_1-H11 using SpyTag-SpyCatcher chemistry^50^ (see Methods). We found that the F domain displaying capsids had very high potency in driving FOXO1 exclusion from the nucleus^50–55^ (Fig.4c-d), activating the AKT pathway (Fig.4d), and stabilizing nascent blood vessels formed from HUVECs^50,56,57^ (Fig.4e). In comparison, 0.17 nM capsid-Fd fusion particles (10 nM equivalent of Fd-protein) elicited a stronger response than that of 100 nM Angiopoetin 1 (Ang1). This designed capsid is also far easier to produce and much more stable than Ang1 and hence these constructs could have practical application to stimulating differentiation and regeneration *ex vivo* and ultimately *in vivo*.

The high surface presentation density enabled by the designed scaffolds provides a route to investigating the effect of packing density on the elicitation of immune responses by nanoparticle based immunogens. As a first step in this direction, we fused trimeric influenza hemagglutinin (HA) fusions onto the N-terminus of I1-capsid RC_I_1-H11 through (GS)5 linkers (Fig.4f). The fusion protein was expressed and secreted from mammalian cells, and clearly forms HA-displaying particles according to SEC and nsEM. Biolayer interferometry (BLI) showed binding of both 5J8 (anti-HA head antibody) and CR9114 (anti-HA stem antibody) to HA capsids (Fig.4g), indicating that the HA remains antigenically intact as displayed on the surface of the capsids.

## Conclusion

The design of 54 and 67 residue proteins that accurately assemble into 60 subunit icosahedra with both internal monomer and overall assembly structure nearly identical to the computational method highlights the power of our top-down RL approach for exploring new protein nanomaterial architectures. The designs are distinct from any previously designed or naturally occurring cages, with smaller subunits, smaller radii, and lower porosities. These structures could not have been built with previous bottom-up approaches, as assembly is fully cooperative and there are no stable cyclic oligomeric substructures. The density of protein chains and termini available for fusion is considerably greater than the most compact previously designed assembly. The potent activation of the Angiopoietin pathway, as well as the robustness observed for display of complex proteins such as HA suggest that these designs could enable a new generation of multivalent cellular receptor agonists and vaccine candidates. The low porosity and high stability could also enable improved encapsulation and delivery for lower molecular weight cargos. More generally, our results highlight the power of reinforcement learning for protein design, which we expect can be increased further by incorporation of policy and value networks^42–44^ to further guide search.

## Methods

### Monte Carlo tree search backbone sampling

The backbone generation method described here uses Monte Carlo tree search (MCTS), an RL algorithm, to choose secondary structure fragments of helices and loops in a decision tree to append to either terminus of a growing protein backbone. With reinforcement based on provided geometric constraints and score functions, iterations through the tree lead to more and more optimal backbone designs as the tree search is guided to better paths.

#### Helix and loop elements

For this method, fragment assembly through the decision tree uses two types of backbone secondary structure elements to select as choices: helices and loops. Helix choices are selected by number of residues of a parametrically-generated straight alpha-helix. This simplification severely limits the size of the structural search space and corresponding decision tree, allowing for more efficient exploration of the space. For example, allowing helix additions in the range of 9-22 residues in length results in only 14 possible choices of helices. Furthermore, simple parametrically-generated alpha helices have been used successfully to solve a wide variety of protein design problems, and so we hypothesized that this simplified sampling strategy would work adequately for many types of problems while avoiding complications arising from more complicated structural elements such as beta-sheets.

Similarly, we sought to use a library of short loop structures for loop choices to increase the likelihood of sampling successful designs. We used a library of approximately 26 thousand loops of 3-5 residues in length extracted from a set of de novo 4 helix bundles designed using the Rosetta BluePrintBDR^58^ and filtered by loop RMSD to the PDB^59^. We chose to bin the loops by structural similarity in order to reduce the number of choices in the decision tree and allow for alternate loop selection as a means to provide slight structural deviation and diversity. Loop choices in the decision tree are therefore the selection of a loop bin followed by the random selection of a loop from within that bin. This strategy required clustering based on parameters describing the helix following loop addition to enable structural similarity in subsequent choices using different loops within the same bin. We used k-means clustering on a set of 7 parameters describing the helix following each loop: 3 for helix translation, 3 for helix direction vector, and 1 for helix phase. We wanted a limited number of bins with a minimum number of loops per bin and a limited range in the distribution of loops per bin in order to avoid sampling biases resulting from over or under populated bins. The loop library was binned into 316 bins representing 316 possible loop choices in the decision tree, where each bin contains between 5 and 392 loops, with 81 loops in a bin on average.

#### Initialization and decision tree choices

To build backbones using this decision tree, a starting point is required for initialization. This starting point is fixed for a given tree, as backbones are built in place so a sequence of choices to sample a given backbone may only be valid in the context of a single starting point. This starting point can be an alpha-helical ‘stub’, placed in space with random translation and rotation, or a pre-existing structure off of which to build. Helix and loop choices from the decision tree are appended to the starting point to build the backbone using the npose package^60^ to align the chosen element to the growing structure. The loops in the loop library have additional short 4-residue helices on either end to allow for accurate alignment to helices. Helix and loop choices may be appended to only a single terminus (N or C) or to both termini as specified by the user. For example, when building off of a randomly placed stub (as used in all cases in this work), the first set of choices in the decision tree will be 14 options for helix lengths between 9 and 22 residues extended off of the C terminus. The following choices in the next level of the tree consist of 732 options, or 316 loop bins each for loop additions off of either terminus. After a loop is chosen, the only allowed options are helix additions from the same terminus, as each loop must be followed by a helix. We found empirically that sampling optimal structures of some number of helices is much more efficient when building from both termini. This is likely due to a reduction of the lever arm effects found when substituting one loop for another similar loop from the same bin, where small deviations can result in much larger propagated structural changes if many subsequent choices are appended to the same terminus. This means that more backbones with slight deviations as well as backbones with similar starting structure choices are able to be sampled.

#### MCTS expansion and evaluation

Random paths through this decision tree of helix and loop choices may lead to viable backbones, but it is computationally intractable to exhaustively explore all paths through the tree and most paths will result in either poor quality backbones that are suboptimal or invalid backbones that violate geometric constraints. With MCTS, paths are initially chosen with uniform random probability, but each iteration informs the choice of subsequent paths to guide the sampling towards better backbones. Upon initialization, all possible choices at each new branch point in the decision tree are assigned the same probability weight. Every choice is made at random, but weights may be adjusted with every iteration to increase their likelihood of selection. These probability weights are changed for each sampled backbone based on geometric constraints that are assessed at every choice as well as score functions that are only assessed for the final structure. Both geometric constraints and score functions must be rapidly calculated to allow for efficient sampling through thousands of iterations of the tree search. As score functions are typically slower to assess and often require a full structure for accuracy, they are only calculated on final structures that pass a specified minimum sequence length and minimum number of helices.

#### Geometric constraints

Geometric constraints are rapidly assessed for each helix and loop choice so that only valid choices are accepted. The first type of geometric constraint is the clash constraint, used for all backbone sampling with this method. Distances are calculated between all new backbone atoms for the potential helix or loop option and all other specified atoms. These specified atoms always include the growing backbone that is being sampled through the tree search, but may also include atoms of other prespecified structures to avoid in space. These other atoms can also include neighboring subunits when sampling symmetric assemblies, as was used for sampling the icosahedral backbones. With every new potential choice, transformation matrices are used to calculate the atoms of all nearby subunits with this additional choice (rather than every subunit to speed up computation), so that symmetric assemblies can be built in place while ensuring no clashes. For all clash checks, all atom distances must be above a specified clash threshold of 2.85 Å.

Simple space bounds may also be provided as geometric constraints, for example limits in Cartesian space where new atoms may not be placed. The diameter of the growing backbone can also be geometrically constrained by calculating distances between all atoms of the growing structure so that no new choice is allowed where any distance exceeds a specified diameter threshold. More complicated volume constraints can be provided through the use of *.obj* files specifying a mesh surface. A grid of points 1Å apart encompassing the entire volume is queried using the pymesh package^61^ to find a winding number for each point. This number specifies whether the point falls within or outside of the volume. All points falling within the volume (winding number >= 1) are added to a hash table specifying all voxels of that volume. All atoms for every potential helix or loop choice are converted to their corresponding voxels, and these voxels are rapidly checked for their presence in the hash table such that only choices where all new atoms are within the specified volume are allowed.

As geometric constraints are assessed for every choice in the tree, only valid choices are allowed. With each valid choice selected, the probability of that choice is upweighted by a fixed value (set to 100), as are the probabilities of every prior choice leading back up the path in the tree. This way, a valid choice is rewarded not only for its initial selection, but also for subsequent valid choices that it enables.

#### Termination cases

Each iteration through the search tree will end at one of a number of termination cases. As only choices passing all geometric constraints are permitted, termination may occur after reaching a set limit for the number of invalid choice attempts at a given branch point. This trial limit serves to constrain the amount of time spent on an iteration, as many paths will lead to points in the tree where few or no possible valid choices exist. The trial limit is 4 at the start of the tree, and increases by an additional value of 3 times the number of helices progressing down the tree, as later branch points will typically have fewer valid choice options and thus require additional attempts to find a valid choice if one exists. Furthermore, it is advantageous to spend compute time on iterations where part of a valid backbone is built already over iterations reaching early dead-end paths. Iterations may also terminate upon reaching preset limits on sequence length and number of helices, which are checked following each choice addition. Upon termination in instances where the last addition was a loop, the loop is removed to avoid lower quality backbones with dangling loops lacking a succeeding helix. Lastly, termination may occur upon picking an additional ‘no choice’, option. This option is introduced as an additional choice at loop branch points after passing a prespecified number of helices, and provides a means of terminating backbones that may already be of good quality that would be worsened with the addition of further choices. The probability for the ‘no choice’, termination is initialized at 0.1 when the minimum number of helices is first reached, then is initialized with an added probability of 0.05 for each helix added past the minimum, as it will typically become a more optimal option. Upon all cases of termination, backbones that exceed the minimum number of helices and an additional preset minimum sequence length are evaluated using specified score functions, and the tree path probability weights corresponding to the backbone are updated.

#### Score functions

Score functions are only evaluated at the end of an iteration on backbones that exceed a minimum number of helices and minimum sequence length, as score functions may be computationally costly and typically only make sense to evaluate on a completed structure that resembles desired backbones. Empirically, we found the MCTS more efficiently produces quality backbones with this strategy than when assessing score functions and upweighting at every step or for backbones beneath minimum helix number and sequence length limits. In testing a wide variety of score functions, we found that the MCTS was always able to enrich for better scoring backbones. We observed that score functions prioritizing computational speed through approximation were more efficient than slower, more accurate score functions, and typically sufficient for producing backbones that still scored optimally using the slower score functions. We also observed that broader, more general score functions allowed the MCTS to produce better backbones, as the method is powerful in finding diverse backbone solutions but will exploit inadequacies in score functions when overly constrained. With these observations in mind, we developed score functions optimizing for speed and generalization.

In this work, we describe three score functions assessing monomer quality. These functions attempt to reward backbones that could potentially be designed into stable monomers. The core formation score uses a cone in front of each residue’s C alpha - C beta vector to calculate the percent of the backbone to classify as ‘core, by sidechain neighbors^62^. The helix packing score uses the same sidechain neighbors calculation to quantify a worst helix, which has the minimum average sidechain neighbors. This helix may be penalized, as if it dangles off of the structure it could lead to an unstable monomer. Lastly, the monomer designability score, repurposed from RPXdock, calculates transforms for all residue pairs and uses them to check a hash table for pair ‘hits’, suggesting sidechain designability^20^. The score function returns the percent of 9mers with hash table hits in the core or core boundary region (classified by sidechain neighbors), indicating the potential designability of the monomer backbone.

We developed two additional score functions for assessing icosahedral assembly quality. These functions attempt to reward assemblies with closed surfaces and good interfaces between subunits. The porosity score rapidly approximates how porous the surface of the icosahedral assembly is by calculating the maximum volume filled across every spherical shell bounded by two atoms. The set of shells is defined by the radial distances of all pairs of atoms in a single subunit, plus an additional carbon Van der Waal (VDW) radius (1.7 Å) for the outer atom and minus the same 1.7 Å for the inner atom. For each of these shells, the approximate volume filled is the sum of all atom sphere volumes divided by the volume of the shell, using the carbon VDW radius once more to calculate all atom sphere volumes. The maximum fraction filled for the set of shells approximates the porosity of the icosahedral assembly. We developed a number of different alternative porosity functions, but found that this particular function was most effective in balancing speed while handling diverse surface closure modes. The porosity score also indirectly enriches for assemblies with better interfaces, as surface closure requires contact between neighboring subunits. The interface designability score further assesses interface quality by using the same atom pair transform hash table as the monomer designability score. The function checks atom pair transforms from one subunit to all neighboring subunits, and returns the total number of hash table hits only if there are hits present across two or more interfaces, thus evaluating the presence and designability of multiple subunit interfaces.

After evaluation, all scores are combined into a single value to reward backbones that are optimal across many or all provided score functions. This is done by passing each score through a sigmoid activation function, with parameters specifying the desired score values and weight relative to other scores. The product of these sigmoid-activated scores is passed through a final sigmoid activation provided it exceeds a minimum threshold, resulting in a final overall score between 0 and 1. This overall score is multiplied by a fixed scalar (set to 5,000) then used to upweight the probability weights of each choice down the decision tree path leading to the final backbone.

#### Upweighting

Probabilities for the path of decision tree choices are upweighted at every step for satisfying geometric constraints and upon termination after evaluating score functions. The relative values of the upweighting amounts and the initialized probability weights determine the extent to which upweighting will affect subsequent iterations through the tree, by increasing the likelihood that a given choice will be randomly selected again. Loop choices have an initialized probability weight of 10, while helix initial probability weights are variable depending on allowed lengths. As there are many more loop options than helix options at each branch point, the starting weights for helices are normalized such that the sum of helix choice weights equals the sum of loop choice weights. This adjustment ensures that a given upweighting amount will increase the probability of a helix or a loop choice equally, as otherwise helix choice diversity would be lost much more rapidly. As discussed earlier, the additional ‘no choice’, termination option is added separately with its own initialization weight, which begins at 10% of the sum of all other choices at a loop branch point and increases by 5% for each additional helix past the minimum helix specification. Geometric constraints are upweighted for a path through the tree by adding a value of 100 to each probability weight, while score functions are upweighted by adding a value of 5,000 multiplied by the calculated overall score, which is between 0 and 1.

To avoid excessive upweighting of single choices leading to a loss of diversity in output structures in later iterations, we implemented a limit to the maximum probability that any single choice may have. At the start of the tree this limit is 0.4, and increases by a factor of 0.025 times the number of choices down the tree up to a maximum value of 0.6. In instances where upweighting would exceed this limit, the probability weight is instead set to a value to equal the probability limit.

#### Optimization

A number of different parameters affect how a MCTS run will proceed and in turn the quality of the resulting backbones, including values describing choice trial attempts, probability weights and upweighting, relative score contributions in sigmoid activations, and total iterations. Although our initial naive choices for these values were able to produce some quality backbones, we found through adjustment we could increase the efficiency of the sampling, producing more acceptable backbones within the same compute time. Further, we sought to balance exploration and exploitation to find settings that produced both diverse and high quality backbones. However, optimization for this model is challenging, as the stochastic nature of the search produces very noisy results and the parameter space is large. Optimal values will also vary across different protein design problems, different initialization points for the same design problem setup, or even different runs using the same initialization. We settled on the values found in the code after many rounds of optimization and several different qualitative and quantitative means of assessing output quality and efficiency, but acknowledge that further optimization is definitely possible.

#### Volume filling production runs

To sample backbones within volumes representing a variety of shapes (supplementary figure 1), we first generated the desired shapes using CAD software. These shapes were converted to volume hash tables to use as geometric constraints. For each shape, we ran thousands of different MCTS runs, each initialized randomly within the volume. We first used 500 test iterations to check that backbone lengths and scores were improving indicating a reasonable initialization point, then ran an additional 50,000 iterations for each passing MCTS run. We permitted helices between 6 and 23 residues in length, and set a minimum of 6 helices and 100 residues of sequence length. For the purpose of filling the desired shapes, we provided only sequence length as a score function, so that the MCTS would prioritize finding paths that fit as many residues in the volume as possible. For supplementary figure 1, we provide for each shape the 5 longest backbones found, as well as our favorite backbone from the same set of 5 most closely resembling the shape.

#### Icosahedral assembly sampling production runs

To sample icosahedral backbones, we used random stub initialization within a sphere of radius 60A. For each MCTS run, we used 500 test iterations to check that the initialization point was reasonable, followed by 10,000 iterations. We permitted helices between 9 and 22 residues in length. We set a minimum of 3 helices, a maximum of 7 helices, a minimum of 50 residues, and a maximum of 80 residues. We set a maximum radial distance of 75A to constrain the size of the assemblies. We found that this search space was constrained enough to efficiently find a large number of diverse and high quality backbones of similar sizes. We used the three monomer score functions and two icosahedral assembly score functions described previously. For each run, we output at most the 5 highest scoring backbones, where all output backbones required a core formation score greater than or equal to 0.2, a helix packing score greater than 2.0, a monomer designability score greater than or equal to 0.9, a porosity score greater than or equal to 0.45, and an interface designability score greater than 17. Each run produced on average fewer than 1 backbone with these strict score thresholds. We repeated the randomly-initialized MCTS icosahedral assembly run millions of times, each with an independent tree explored through 10,000 iterations, to produce a library of approximately 1 million icosahedral backbones. Tree search iterations took on average 0.045 seconds, with each run taking on average 7.5 minutes. We estimate this library took approximately 1.5 million CPU hours to generate.

### Sequence length and porosity analysis of native, bottom-up, and top-down icosahedra

To perform the sequence length and porosity analysis in Figure 1d, we first downloaded all icosahedral homo 60-mers from the PDB (as of August 1, 2022). All partial and non-native structures were removed, as well as the much longer sequence length PDB 3J3I for graph clarity, leaving a library of 240 icosahedra. A total of 3 icosahedral homo 60-mer “bottom-up” structures were obtained from databases at the IPD. A random selection of 25 MCTS-generated designs were selected for the “top-down” structures from the set of designs screened experimentally. Amino acid sequence length for a single subunit was determined from the structure files. A ray tracing method was utilized to find a measure of porosity for each structure, in which 10,000 rays with random direction were traced from the origin to the outside of the icosahedron. The fraction of rays that leave the capsid without coming within 0.5 Å of an amino acid atom is reported as the porosity. The choice of 10,000 random rays was benchmarked for 10 random native icosahedra against a larger more accurate run of 100,000 random rays, and found to be > 95% accurate in all 10 cases.

### Protein BiRNN model for sequence profile prediction

The trRosetta^63^ training set was used to train a deep neural network (ProteinBiRNN) for the prediction of each amino acid probability at each residue position given solely the backbone structure. The architecture of the deep neural network consists of several convolutional layers, a bidirectional recurrent neural network and a residue-level attention layer. All the input features are directly derived from the backbone structure and calculated on-the-fly. The 1D features include: 1) per-residue torsion angle phi, psi and omega; one hot encoded secondary structure type (H, L, E); and number of sidechain neighbors). The 2D features include: 1) inter-residue distance; relative orientation geometries (Supplementary Fig.4). The network predicts the amino acid identity of each residue and we used categorical cross-entropy to measure the loss for the training. All the trainable parameters restrained by the L2 penalty and dropout keeping probability 90% is used.

### Sequence design, evaluation, and selection

The computational sequence design pipeline contains multiple modules (Supplementary Fig.3). Position-specific scoring matrices (PSSM) were generated with the ProteinBiRNN model. Using tertiary structures as input, either a monomer or a multi-chain fragment or the entire capsid, the ProteinBiRNN predicts sequence probability distribution matrices for each residue position, based on its local backbone neighborhood environment. For each MCTS-generated capsid backbone, two ProteinBiRNN profile predictions were calculated for both monomer fold (pssm_chA) and asymmetric subunit that contain all interfaces (pssm_int), respectively.

To reduce unnecessary calculations, a fast pre-design step (< 2 min per CPU) was first performed to examine whether a monomer building block would preferentially fold into the designed backbone configuration. Residue type constraints (pssm_chA) were set by using the FavorSequenceProfile mover during Rosetta Fastdesign. Designed monomer scaffolds with unfavorable Rosetta score/residue (> -2.7) and AlphaFold metrics (pLDDT < 80, RMSD > 1.5 Å) were excluded from the pool for downstream studies. For example, in a test set of around 80,000 MCTS-generated backbones, about 30 % of designs passed the above filtering criteria (Supplementary Fig.5).

Next, an interface screening stage of symmetric RosettaDesign calculation was performed for all capsids to design contacting assembly interfaces and protein cores in a single step, as guided by pssm_int. We applied a Rosetta FastDesign protocol, similar to the one described in a recent publication^45^, to activate between side-chain rotamer optimization and gradient-descent-based energy minimization. Symmetric Rosetta backbone relaxation and small random perturbation to local backbone positions were observed to improve upon sequence encoding and interface shape complementarity, as compared to conventional cage design protocols with fixed input backbones^10^. Computational metrics of the final design models were calculated using Rosetta, which includes ddG, shape complementarity and interface buried SASA, contact molecular surface, among others, for design selection (Supplementary Fig.9). AF metrics suggest the sequence encoding obtained by ProteinBiRNN PSSM-guided Rosettadesign outperforms conventional layer design (Supplementary Fig.6). All the script and flag files to run the programs are provided in the Supplementary Information. With backbone relaxation, a typical symmetric capsid design calculation takes about 3.5 hours on average to finish on a CPU.

To evaluate the interface quality of designed capsids, symmetric interfaces between two neighboring chains (C2,C3, and C5) were extracted and analyzed independently (Supplementary Fig.7). To ensure capsid formation, we hypothesized that at least two relatively strong interfaces need to exist, where a first one is needed for homo-oligomeric formation, and a second one closes the capisd. Inspired by previous studies^10^, we set the cutoff for relatively strong interface based on its calculated Rosetta metrics, including binding energy (ddG < −20) and contact molecular surface (cms >180) with appreciable hydrophobic side chain packing (cms_apolar> 120).

During the final sequence optimization stage, Rosetta and ProteinMPNN^49^ were used to design the selected capsid set with good backbone dockings (≥ 2 strong interfaces). An iterative backbone optimization process was cycled three times during the final sequence optimization step to improve interface shape complementarity and designed backbone robustness. For each design round, the Rosetta relaxed design model and its corresponding AF monomer prediction (from previous round) were TMaligned^64^ back to the starting position and used as input for capsid design using Rosetta or ProteinMPNN (Supplementary Fig.8). To design capsid sequences with the ProteinMPNN module, the tertiary structure of a full capsid was used as input while all 60 sequences were tied to be identical to enforce icosahedral symmetry. A typical proteinMPNN design calculation takes about 5 mins on a GPU for a homo-60 mer capsid with ~ 70 residues per monomer, which is significantly faster than conventional Rosetta Fastdesign. To generate pdb models for the MPNN output sequences, the output sequences were grafted onto the input backbone using Rosetta SimpleThreadingMover^65^ followed by symmetric relaxation. Finally, all final capsid designs were evaluated by Rosetta (monomer and interface metrics) and Alphafold (monomer pLDDT and RMSD) for design selection. Detailed filter setup can be found in the Supplementary Information.

### Protein expression and purification

For small scale screening, synthetic genes encoding capsid sequences were optimized for E. coli expression and purchased from IDT (Integrated DNA Technologies) as linear eblock DNA in 96-well format. We performed Golden gate assemblies following the standard protocol (New England Biolab) to clone each synthetic gene into a pET29b vector with a hexahistidine affinity tag. Plasmids were then transformed into BL21* (DE3) (Invitrogen) E. coli competent cells and grew overnight in 1 mL of Studier autoinduction media. The cells were harvested by centrifugation, resuspended and lysed in 200 μL Bugbuster protein extraction reagent (Millipore). Following another centrifugation, the supernatant was purified by Ni^2+^ immobilized metal affinity chromatography (IMAC) with Ni-NTA charged magnetic beads (Qiagen). Magnetic beads with bound cell lysate were washed with 200 μL of washing buffer for three times (150 mM NaCl, 25 mM Tris pH 8.0, 30 mM imidazole) and eluted with 35 μL of elution buffer (150 mM NaCl, 25 mM Tris pH 8.0, 500 mM imidazole). Soluble fractions were analyzed by SDS-PAGE to check for a single intense band at expected molecular weights. The selected IMAC elution of soluble designs were diluted to appropriate concentration with a buffer (1:20, 150 mM NaCl, 25 mM Tris pH = 8.0) and analyzed by negative-stain electron microscopy (see below).

For large scale protein expression, plasmids were transformed into BL21* (DE3) E. coli competent cells. Single colonies from agar plate with 100 mg/L kanamycin were inoculated in 50 mL of studier autoinduction media, and the expression continued at 37 °C for over 24 hours. The cells were harvested by centrifugation, resuspended in 150 mM NaCl, 25 mM Tris pH 8.0, and lysed by 20 mL Bugbuster protein extraction buffer (Millipore, 50 mL Bugbuster lysis buffer contains - 49 mL Bugbuster reagent, 0.75 mL 2M imidazole, 1 protease inhibitor tablet, 50 mg lysozyme). Following another centrifugation, the supernatant was purified by 1 mL Ni^2+^ IMAC with Ni-NTA Superflow resins (Qiagen). Resins with bound cell lysate were washed with 10 mL (bed volume 1 mL) of washing buffer (same as above) for two times and eluted with 3 mL of elution buffer (same as above). To improve colloidal stability of the capsid protein, glycine was added to the IMAC elution to reach a final concentration of 100 mM, which was subsequently concentrated and purified via size exclusion chromatography (SEC) in 25 mM pH 8.0 Tris buffer (150 mM

NaCl) on a Superose 6 Increase 10/300 gel filtration column (Cytiva). The resulting samples were generally > 95 % homogeneous on SDS-PAGE gels. SEC-purified designs were concentrated by 10K concentrators (Amicon) and quantified by UV absorbance at 280 nm and stocked at 4 °C.

The HA mini capsid gene was composed of the A/Michigan/57/2015 H1N1 HA ectodomain with the Y98F mutation1 (PMID: 24501410) fused via a 5(GS) linker to the RC_I_1 capsid with an N-terminal hexahistidine affinity tag. This gene was codon optimized for human cell expression and made in the CMV/R mammalian expression vector by Genscript. Transient transfection into HEK293F cells was carried out using PEI MAX. After 4 days of expression, mammalian cell supernatants were clarified via centrifugation and filtration. IMAC was used to initially purify HA capsids from cell supernatant by adding in 1 ml of Ni2+ sepharose excel resin per 100 ml supernatant, along with 5 ml of 1 M Tris, pH 8.0 and 7 ml of 5 M NaCl. This batch binding was left to proceed for 30 min at room temperature with shaking. Resin was then collected in a gravity column, washed with 5 column volumes of 50 mM Tris, pH 8.0, 500 mM NaCl, 20 mM imidazole, and his-tagged HA capsids were eluted using 50 mM Tris, pH 8.0, 500 mM NaCl, 300 mM imidazole. Purification by SEC was then carried out on a Superose 6 Increase 10/300 gel filtration column equilibrated in 25 mM Tris, pH 8.0, 150 mM NaCl.

### Bio-layer interferometry (BLI)

BLI was carried out using an Octet Red 96 system, at 25°C with 1000 rpm shaking. Antibodies were diluted to 10 ug/ml in kinetics buffer (PBS with 0.5% serum bovine albumin and 0.01% Tween) and then loaded onto protein A tips for 200 s. HA mini capsid was diluted to 500 nM in kinetics buffer and its association was measured for 200 s, followed by dissociation for 200 s in kinetics buffer alone.

### Transmission negative-stain electron microscopy (nsEM)

Selected IMAC elutions and cage fractions from SEC traces were diluted to about 0.5 μM (monomeric component concentration) for negative-stain EM characterization. A drop of 5 μL sample was applied on negatively glow discharged, formvar/carbon supported 400-mesh copper grids (Ted Pella, Inc.) for 1 min. The grid was blotted and stained with 5 μL of uranyl formate, blotted again, and stained with another 5 μL of uranyl formate for 60 s before final blotting. 2% uranyl formate was used for all samples.

The screening and data collection was performed on a 120kV Talos L120C transmission electron microscope (Thermo Scientific) with a BM-Ceta camera using EPU 2.0. All nsEM datasets were processed by the CryoSparc software^66^. Micrographs were imported into the CryoSparc web server and the contrast transfer function (CTF) was corrected. All the picked particles were 2D classified for 20 iterations into 50 classes. Particles from selected classes were used for building the ab-initio initial model. The initial 3D model was homogeneously refined using C1 and the corresponding I symmetry.

### Circular dichroism (CD) experiments

To study the secondary structure and thermodynamics of the designed capsid proteins, CD measurements were performed with an JASCO 1500. The 200 to 195 nm wavelength scans were measured at every 10 °C intervals from 25 to 95 °C. For each measurement, a 1-mm path length cuvette was loaded with protein concentrations at 0.2 mg/mL in TBS buffer, as measured by Nanodrop at 280 nm using predicted extinction coefficients.

### CryoEM sample preparation

To prepare cryoEM sample grids for the capsids, 3 μL of 0.5 - 1.0 mg/mL of capsid proteins in 150 mM NaCl, 25 mM Tris (pH = 8.0), 100 mM Glycine was applied to glow-discharged Quantifoil R 2/2 300 mesh copper grids overlaid with a thin layer of carbon. Vitrification was performed on a Mark IV Vitrobot with a wait time of either 5 or 7.5 seconds, a blot time of 0.5 seconds, and a blot force of either 0 or -1 before being immediately plunged frozen into liquid ethane. The sample grids were clipped following standard protocols before being loaded into the microscope for imaging.

### CryoEM data collection

Capsid data collection was performed automatically using either Leginon^67^ or SerialEM to control either a ThermoFisher Titan Krios 300 kV equipped with a standalone K3 Summit direct electron detector with an energy filter, or a ThermoFisher Glacios 200 kV equipped with a standalone K2 Summit direct electron detector^68^. All three capsids were collected using counting mode with random defocus ranges spanned between -0.7 and -2.0 μm using image shift, with one-shot per hole on a Glacios for RC_1-H11, and 6 shots per hole on a Krios for RC_I_1 and RC_I_2. 3,888, 12,825, and 1,315 movies were collected with a pixel size of 0.84 Å, 0.84 Å, and 1.16 Å for RC_I_1, RC_I_2, and RC_I_1-H11 capsids, respectively. A total dose of ~60 e^-^/Å^2^ was applied to RC_I_1 and RC_I_2 on a Titan Krios, and ~50 e^-^/Å^2^ for RC_1_I-H11 on a Glacios.

### CryoEM data processing

All data processing was carried out in CryoSPARC and CryoSPARC Live^66^. Alignment of movie frames was performed using Patch Motion with an estimated B-factor of 500 Å^2^, with a maximum alignment resolution set to 3. Defocus and astigmatism values were estimated using Patch CTF with default parameters. RC_I_1 particles were initially picked in a reference-free manner using Blob Picker and extracted with a box size of 320 pixels. This was followed by a round of 2D classification and subsequent template-picking using the best 2D class averages low-pass filtered to 20 Å, for a total of 805,450 picked particles. For RC_I_1-H-11, 491,297 particles were picked with Template Picker using templates from the RC_I_1 dataset, and were extracted with a box size of 320 pixels. For RC_I_2, 769,399 particles were ultimately picked and extracted with a box size of 320 pixels after a round of reference-free picking using Blob Picker and a subsequent Template Picker job using templates derived from 2D class averages which were low-pass filtered to 20 Å. Rounds of reference-free 2D classification were next performed in CryoSPARC with a maximum alignment resolution of 6 Å for each dataset. The best classes for each sample that revealed clearly visible secondary-structural elements were used for 3D ab initio determination using the C1 symmetry operator. RC_I_1 and RC_I_1-H-11 datasets were next corrected for local particle motion followed by 3D ab initio using C1 and I symmetry, respectively. This was followed by a 3D refinement with I symmetry for RC_I_1 for a final global resolution estimate of 2.5 Å. For RC-I_1-H-11, ab initio was followed by Non-Uniform Refinement and Local Refinement for a final global resolution estimate of 3.0 Å. For RC_I_2, data was processed entirely in CryoSPARC Live during collection. After selection of the best 2D class averages following template picking, an ab-initio job was run using C1 symmetry on 100,000 particles, and a final refinement performed with I symmetry applied using 283,552 of the best particles from 2D classification, yielding a global resolution estimate of 2.94 Å. Local resolution estimates were determined in CryoSPARC using an FSC threshold of 0.143. 3D maps for the half maps, final unsharpened maps, and the final sharpened maps for each capsids RC_I_1, RC_I_1-H11, and RC_I_2 were deposited in the EMDB under accession number EMD-xxxx, EMD-xxxx, and EMD-xxxx, respectively.

### CryoEM model building and validation

The *de novo* predicted design models for each capsid (reported here) were used as initial references for building the final cryoEM structures. The models were manually edited and trimmed using Coot^69,70^. We further refined each structure in Rosetta using density-guided protocols^71^. EM density-guided molecular dynamics simulations were next performed using Interactive Structure Optimization by Local Direct Exploration (ISOLDE), with manual local inspection and guided correction of rotamers and clashes throughout simulated iterations. ISOLDE runs were performed at a simulated 25 Kelvin, with a round of Rosetta density-guided relaxation performed afterward. This process was repeated iteratively until convergence and high agreement with the map was achieved. Multiple rounds of relaxation and minimization were performed on the complete capsids, followed by human inspection for errors after each step. Throughout this process, we applied strict non-crystallographic symmetry constraints in Rosetta^71^. Phenix real-space refinement was subsequently performed as a final step before the final model quality was analyzed using Molprobity^72^ and EM ringer^73^. Figures were generated using either UCSF Chimera^74^ or UCSF ChimeraX^75^. The final structures for RC_I_1, RC_I_2, and RC_I_1-H11 were deposited under PDB accession numbers XXXX, XXXX, and XXXX, respectively.

### Cell culture

Human Umbilical Vein Endothelial Cells (HUVECs) were acquired from Lonza (C2519AS). Cells were grown on 0.1% gelatin-coated (Sigma, G1890-100G) 35-mm cell culture dish in EGM2 media described previously^59^. HUVECs were expanded and serially passaged to reach passage 7 before experiments.

### Cap-Fd conjugation

F-domain fused with SpyTag were incubated with RC_I_1-H11 capsid fused with SpyCatcher for 4 hours at room temperature on nultation. Samples were analyzed on a SDS–PAGE gel to confirm at least 90% conjugation was reached (Supplementary Fig.16).

### pAKT titration

Passage 7 HUVECs at 80% confluency were rinsed with 1× PBS (Gibco, 10010023) twice. The cells were then starved in DMEM low glucose 1 g/l (Gibco, 11885-084) serum-free media per plate for 16 hours. At 16-h timepoint, cells will be treated with Cap-Fd (1000-0.1 nM), Fd-st (100 nM), Ang1 (100 nM), or PBS for 15 minutes at 37°C. After treatment, the media was aspirated, and cells were washed once with 1 × PBS before harvesting protein for immunoblotting (Supplementary Fig.17).

### Immunoblotting

Cells were lysed with 130 μl of previously described lysis buffer^50^. Cell lysates were collected in a fresh Eppendorf tube with 4× Laemmle Sample buffer (Bio-Rad, 1610747) containing 10% betamercaptoethanol (Sigma-Aldrich, M7522-100) added and then heated at 95°C for 10 min. 30 μl of protein sample per well was loaded and separated on a 4–10% SDS–PAGE gel for 30 min at 250 Volt using running buffer (7.2 g glycine, 1.5 g Tris-base, and 0.5g SDS diluted in 1L DI water). The proteins were then transferred on a nitrocellulose membrane for 12 min using transfer butter (7.3 g Tris-Base, 3.6 g glycine, and 0.46 g SDS in 1L DI water). Post-transfer, the membrane was blocked in 5% bovine serum albumin for 1 hour. Then the membrane was probed with primary antibodies overnight: pAkt-S473 (Cell Signaling, 9271S) at 1:2,000, b-Actin (Cell Signaling, 3700S) at 1:10,000, and S6 (Cell Signaling, 2217S) at 1:1000. Membranes with primary antibodies were incubated at 4°C. overnight on a rocker. Then the membranes were washed with TBST buffer (2.4 g Tri-HCl, 8 g NaCl, and 1 mL Tween-20 diluted in 1L water with pH adjusted to 7.4) 3 times at 5-min intervals. Following washes, the pAKT membrane was blocked in 5% milk at room temperature for 1 hour and then incubated in the respective HRP-conjugated secondary antibody (1:2,000) prepared in 5% milk for 1 hour. All other primary antibodies were removed and washed 3 times before adding secondary antibodies at 1:10,000 for 1-hour incubation. After 1 hour, membranes were washed with TBST (3 times, 5 min of interval) and developed using Chemiluminescence developer and imaged using Bio-Rad ChemiDoc Imager. Data were quantified using the ImageJ software to analyze band intensity. pAKT band intensity was divided by band intensity of actin or s6. All signaling levels are normalized to Cap-Fd at 10 nM signal levels as an internal positive control. EC50 is calculated using FindEC anything in Prism, GraphPad.

### Immunofluorescence staining of FOXO1

FOXO1 analysis was done as described before^50^. Briefly, passage 7 HUVECs were seeded on glass coverslips coated with 0.1% gelatin and cultured until confluency. Once confluent, cells were starved for 16 hours in low glucose DMEM (1 g/l D-glucose). Then, cells were stimulated Cap-Fd, Fd-st, or Ang1 at 100 nM of F-domain concentration for 15 min before fixing with 4% PFA (EMS, 15710) for 15 min. The fixed cells were washed three times at 5 min each before blocking for 1 hour with 3% BSA and 0.1% Triton in PBS while on nutation. The cells were incubated with FOXO1 (Cell Signaling, 2880) antibody diluted at 1:100 in blocking agent overnight. After the primary antibody, the cells were washed 3 times at 5 min each with PBS while on nutation. The cells were then incubated with secondary antibodies at 1:200 for 1 hour and 20 min at 37°C. After secondary antibodies, cells were washed for three times at 10 min each with PBS on nutation. Coverslips were sealed using VECTASHIELD plus DAPI (Vector laboratories, H-2000-2) upside-down on glass slides for analysis in confocal (Leica).

### Tube formation assay

Tube formation was assessed usingpreviously described protocols^50,56^. Briefly, a 24-well plate was precoated with 150 uL of 100% Matrigel for 30 minutes at room temperature before cell seeding. Passage 6 HUVECs were seeded at 1.5 × 10^5^ cells/350uL density suspended in media (DMEM low glucose + 0.5% FBS ± 100nM of Cap-Fd). 24 hours after cell seeding, the old media is aspirated and replaced with fresh media without any treatment. The cells continue to be incubated for up to 72 hours. Capillary-like structures were observed, and 20 randomly selected microscopic fields were photographed under Nikon Eclipse Ti scope. Images were analyzed to quantify the number of nodes, meshes, and tubes in each image using the Angiogenesis Analyzer plug-in in ImageJ. Vascular stability was calculated by the average number of nodes, meshes, and tubes. Data were normalized to PBS vehicle as fold change.

## Acknowledgements

We thank F. Dimaio, J. Dauparas, B. Coventry, and T. Huddy for help with computational design and discussion; R. Kibler, L. Miles, and N. Ennist for experimental help; S. Dickinson and J. Quipse for help maintaining and operating the electron microscopes used. This work was conducted at the Advanced Light Source (ALS), a national user facility operated by Lawrence Berkeley National Laboratory on behalf of the Department of Energy, Office of Basic Energy Sciences, through the Integrated Diffraction Analysis Technologies (IDAT) program, supported by DOE Office of Biological and Environmental Research. Additional support comes from the National Institute of Health project ALS-ENABLE (P30 GM124169) and a High-End Instrumentation Grant S10OD018483. This work was supported by National Institute on Aging grant 1U19AG065156-01 (to I.L. and D.B.); Amgen (to S.W.), The Audacious project at the institute for Protein Design (to A.J.B., Z.L., L.C., H.R.B., Y.T.Z., D.B., and N.K.); Novo Nordisk Foundation Grant NNF170C0030446 (to C.N.); the Open Philanthropy Project Universal Flu Vaccine and Improving Rosetta Design (to A.D., N.K., and D.B.); a Microsoft gift (to M.B.).

## Author contributions

I.L. and S.W. contributed equally, and the author order was chosen arbitrarily; citations on CVs etc. will be adjusted accordingly. S.W., I.L., C.N., and D.B. conceptualized the research. I.L. developed the RL backbone generation method. S.W. and C.N. developed the sequence design pipeline. L.C. and M.B. developed the DL sequence profile prediction method. S.W. designed the original capsids and performed the screening, expression and characterization experiments. A.J.B. designed cryoEM experiments and optimization of sample purification conditions, performed the initial cryoEM screening experiments, and optimized the cryoEM freezing conditions. A.J.B, S.W., Z.L. prepared additional cryoEM grids and collected cryoEM data. A.J.B. processed the cryoEM data, built, and solved the structures for each designed capsid. S.W. and A.D. designed and characterized the fusion capsids. A.D. and N.K. designed and performed the immunization studies for the HA-capsid fusions. Y.T.Z and H.R.B. designed and performed the cell signaling assays. All authors analyzed data. D.B. supervised research. S.W., I.L. and D.B. wrote the manuscript with the input from the other authors. All authors revised the manuscript.

## Supplementary Information

This file contains a Supplementary Discussion, Supplementary Table 1-3 and Supplementary Figs 1–17.

